# Pandora: 4-D white matter bundle population-based atlases derived from diffusion MRI fiber tractography

**DOI:** 10.1101/2020.06.12.148999

**Authors:** Colin B Hansen, Qi Yang, Ilwoo Lyu, Francois Rheault, Cailey Kerley, Bramsh Qamar Chandio, Shreyas Fadnavis, Owen Williams, Andrea T. Shafer, Susan M. Resnick, David H. Zald, Laurie Cutting, Warren D Taylor, Brian Boyd, Eleftherios Garyfallidis, Adam W Anderson, Maxime Descoteaux, Bennett A Landman, Kurt G Schilling

## Abstract

Brain atlases have proven to be valuable neuroscience tools for localizing regions of interest and performing statistical inferences on populations. Although many human brain atlases exist, most do not contain information about white matter structures, often neglecting them completely or labelling all white matter as a single homogenous substrate. While few white matter atlases do exist based on diffusion MRI fiber tractography, they are often limited to descriptions of white matter as spatially separate “regions” rather than as white matter “bundles” or fascicles, which are well-known to overlap throughout the brain. Additional limitations include small sample sizes, few white matter pathways, and the use of outdated diffusion models and techniques. Here, we present a new population-based collection of white matter atlases represented in both volumetric and surface coordinates in a standard space. These atlases are based on 2443 subjects, and include 216 white matter bundles derived from 6 different state-of-the-art tractography techniques. This atlas is freely available and will be a useful resource for parcellation and segmentation.

## 1. Background & Summary

The creation and application of medical image-based brain atlases is widespread in neuroanatomy and neuroscience research. Atlases have proven to be a valuable tool to enable studies on individual subjects and facilitate inferences and comparisons of different populations, leading to insights into development, cognition, and disease[1–3]. Through the process of spatial normalization, images can be aligned with atlases to facilitate comparisons of brains across subjects, time, or experimental conditions. Additionally, atlases can be used for label propagation, where anatomical labels are propagated from the atlas to new data in order to identify a priori regions of interest. With these applications in mind, a number of human brain atlases have been created (**Figure 1**), with variations in the number of labels, the regions of the brain that are delineated, the methods used to generate labels, and the population or individuals used to create the atlas (for a review of the existing atlases and their standardization, see recent work by Myers et al.[4]).

**Figure 1.**
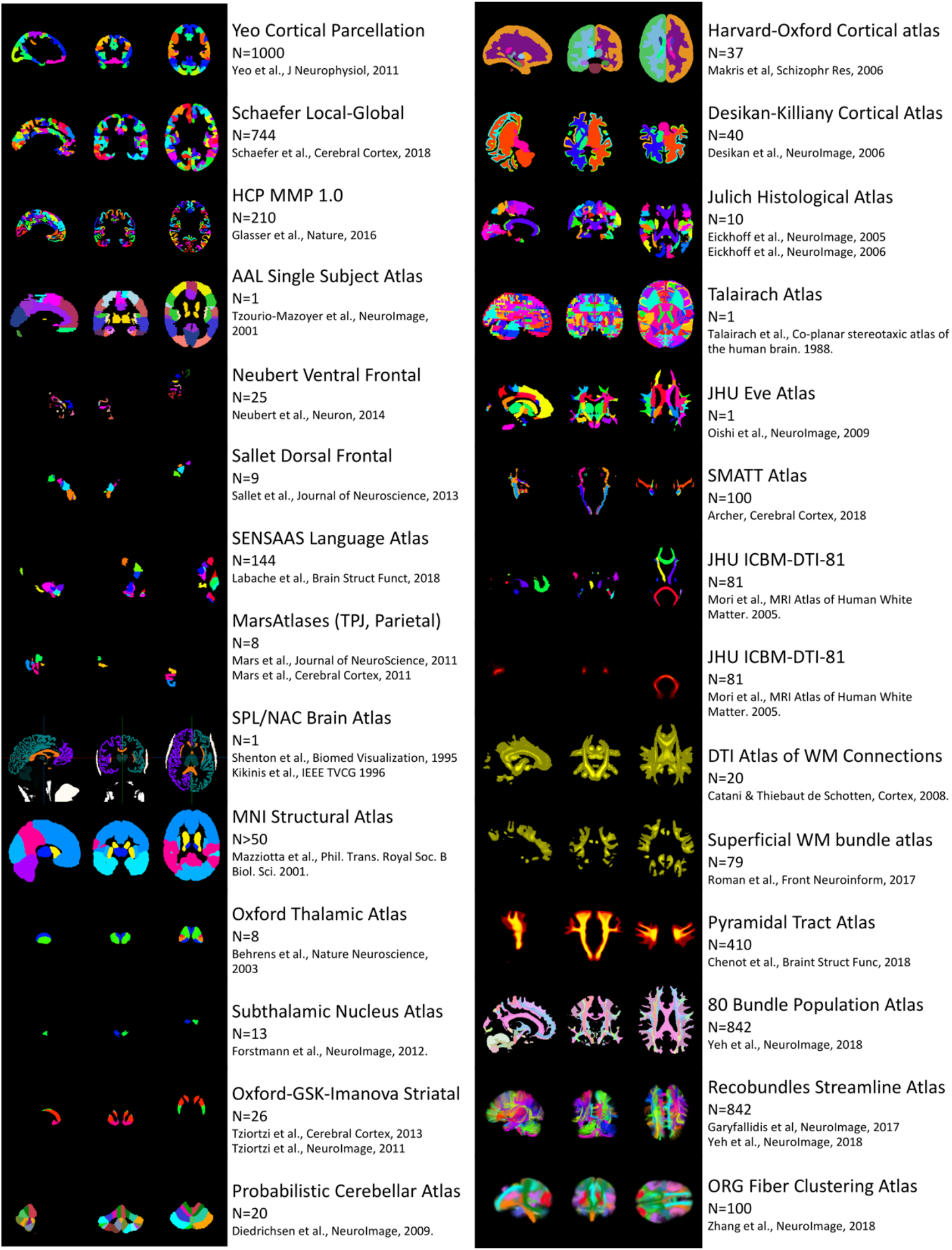
Comparison of types of human brain atlases and regions present in each. Visualizations were made using FSLview tri-planar view for volumetric atlases and using MI-brain 3D-view for streamline atlases. Note that because atlases are in different spaces, visualized slices, anatomy, and orientation is not guaranteed to be the same across atlases. Note that this figure is not exhaustive, and is only representative of the types of atlases and the information they contain. Figure inspired by work on standardizing gray matter parcellations (Figure 1 of Myers et al.^4^).

Despite the wide variety of human brain atlases available to the research community, there is a distinct lack of resources available to describe the white matter of the brain. For example, most atlases emphasize cortical or sub-cortical gray matter, and do not contain a label for white matter [5–24] or only label white matter as a single homogenous structure, or simply separate into the “cerebral white matter” of the left and right hemispheres[25, 26].

Some atlases do indeed include labels for white matter. However, in many cases these labels are for “regions” of the white matter rather than labels for specific white matter bundles[27–32]. For example, an atlas may contain a label for the “anterior limb of the internal capsule” or “corona radiata” which are descriptions of regions through which several white matter bundles are known to pass. While these regions are certainly scientifically useful, the white matter pathways themselves would be more informative for network neuroscience investigations or applications where white matter structure, connectivity, and location are paramount. Additionally, regional labels do not overlap, whereas the fiber bundles of the brain are known to be organized as a complex mixture of structures, overlapping to various degrees.

To overcome these limitations, several atlases have been created using diffusion MRI fiber tractography, a technique which allows the investigator to perform a “virtual dissection” of various white matter bundles of the brain. Examples include population-based atlases of association and projection pathways[33–36], atlases of the superficial U-fibers connecting adjacent gyri[37, 38], and atlases created from tractography on diffusion data averaged over large population cohorts[34, 39, 40]. In particular, several atlases have been made with a focus on a single pathway or a set of pathways with functional relevance, for example the pyramidal tract[41], the sensorimotor tracts[42], or lobular-specific connections[36, 43, 44]. Existing tractography-based atlases, however, typically suffer from one or more limitations: (1) small population sample sizes, (2) restriction to very few white matter pathways, and (3) the use of out-dated modeling for tractography (specifically the use of diffusion tensor imaging which is associated with a number of biases and pitfalls). Further, it is not clear whether the same pathway defined using one atlas results in the same structure when compared to another atlas due to differences in the procedures utilized to define and dissect the bundle under investigation. A final type of atlas, streamline-based atlases[38, 39, 45–47] have become popular in recent years. These are composed of millions of streamlines and can be used as a resource to cluster sets of streamlines on new datasets, thus they nicely complement the use and application of volumetric atlases when diffusion MRI is available.

In this work, we introduce the Pandora^*^ white matter bundle atlas. The Pandora atlas is actually a collection of 4-dimensional population-based atlases represented in both volumetric and surface coordinates in a standard space. Importantly, the atlases are based on a large number of subjects, and are created from multiple state-of-the-art tractography and dissection techniques, resulting in a sizable number of (possibly overlapping) white matter labels. In the following, we describe the creation of these atlases, the data records of the files and their formats, and validate the use of multiple subject populations and multiple tractography methodologies. The Pandora atlas is freely available (https://www.nitrc.org/projects/pandora_atlas; https://github.com/MASILab/Pandora-WhiteMatterAtlas) and will be a useful resource for parcellation and segmentation.

## 2. Methods

**Figure 2** presents an overview of the pipeline and methodology used to create these atlases. Briefly, we retrieved and organized data from 3 large repositories (**Figure 2, Data**). For each subject, we performed six different automated methods of tractography and subsequent white matter dissection (**Figure 2, Subject-level processing: tractography**), and registered all data to a standard volumetric space (**Figure 2, Subject-level processing: registration**). Next, a probabilistic map was created separately for each white matter bundle in standard space in order to create the volumetric atlases (**Figure 2, Volumetric atlas creation**). Finally, a surface mesh of the boundary between white and gray matter was created, and the volumetric maps were used to assign probabilities along this surface to create the surface-intersection atlases (**Figure 2, Surface Atlas creation**). In addition to making the atlases available, all methods are also available as source code (https://github.com/MASILab/Pandora-Methods).

**Figure 2.**
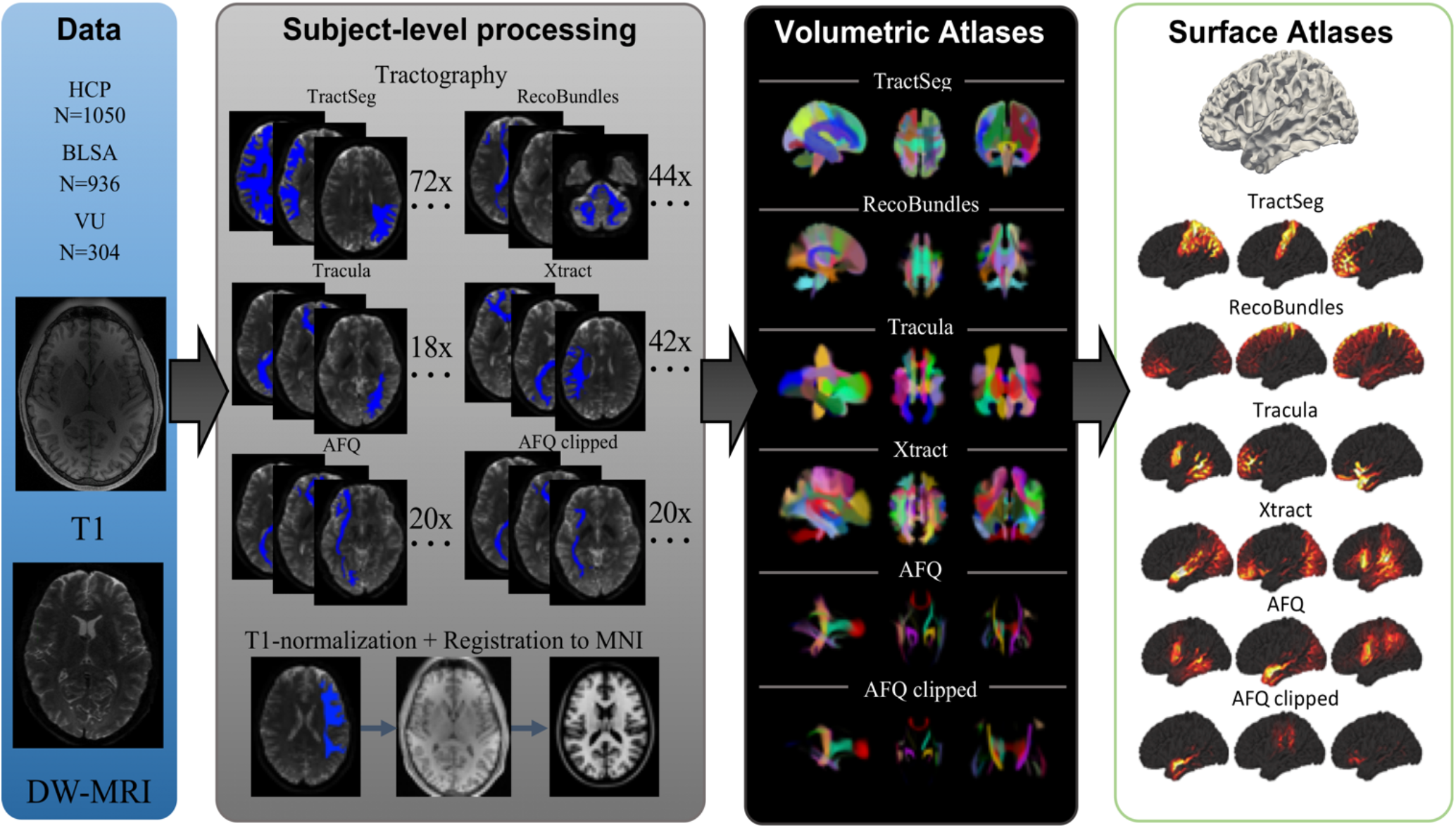
Experimental workflow and generation of Pandora atlases. Data from three repositories (HCP, BLSA, and VU) were curated. Subject-level processing includes tractography and registration to MNI space. Volumetric atlases for each set of bundle definitions is created by population-averaging in standard space. Point clouds are displayed which allow qualitative visualization of probability densities of a number of fiber pathways. Finally, surface atlases are created by assigning indices to the vertices of the MNI template white matter/gray matter boundary.

### 2.1 Data

We used de-identified images from the Baltimore Longitudinal Study of Aging (BLSA), Human Connectome Project (HCP) S1200 release, and Vanderbilt University (**Figure 2, Data**). The BLSA is a long-running study of human aging in community-dwelling volunteers and is conducted by the Intramural Research Program of the National Institute on Aging, NIH. Cognitively normal BLSA participants with diffusion MRI data were included in the present study, using only one scan per participant, even if multiple follow-ups were available. HCP data are freely available and unrestricted for non-commercial research purposes, and are composed of healthy young adults. This study accessed only de-identified participant information. All datasets from Vanderbilt University were acquired as part of a shared database for MRI data gathered from healthy volunteers. A summary of the data is given in **Table 1**, including number of subjects, age, sex, and handedness. All human datasets were acquired under research protocols approved by the local Institutional Review Boards.

**Table 1.** Meta-data information. Note that several inputs are not provided due to confidentiality and data release agreements.

|  | HCP | BLSA | VU |
| --- | --- | --- | --- |
| **Subjects** | 1060 | 963 | 303 |
| **Age** | 28.8±3.5 | 66.2±14.82 | 29.7±11.5 |
| **Age Range** | [22 35] | [22.4 95.1] | [18 75] |
| **Handedness** | N/A | 86L; 843R; 35N/A | 30L; 270R; 3N/A |
| **Sex** | 488M; 572F | 431M; 532F | 134M; 169F |

All datasets included a T1-weighted image, as well as a set of diffusion-weighted images (DWIs). Briefly, the BLSA acquisition (Philips 3T Achieva) included T1-weighted images acquired using an MPRAGE sequence (TE = 3.1 ms, TR = 6.8 ms, slice thickness = 1.2 mm, number of Slices = 170, flip angle = 8 deg, FOV = 256×240mm, acquisition matrix = 256×240, reconstruction matrix = 256×256, reconstructed voxel size = 1×1mm). Diffusion-weighted images were acquired using a single-shot EPI sequence, and consisted of a single b-value (b = 700 s/mm^2^), with 33 volumes (1 b0 + 32 DWIs) acquired axially (TE = 75 ms, TR = 6801 ms, slice thickness = 2.2 mm, number of slices = 65, flip angle = 90 degrees, FOV = 212*212, acquisition matrix = 96*95, reconstruction matrix = 256*256, reconstructed voxel size = 0.83×0.83 mm). HCP acquisition (custom 3T Siemens Skyra) included T1-weighted images acquired using a 3D MPRAGE sequence (TE = 2.1 ms, TR = 2400 ms, slice thickness = 0.7 mm, flip angle = 8 deg, FOV = 224×224mm, acquisition, voxel size = 0.7×0.7mm). Diffusion images were acquired using a single-shot EPI sequence, and consisted of three b-values (b = 1000, 2000, and 3000 s/mm^2^), with 90 directions (and 6 b=0 s/mm^2^) per shell (TE = 89.5 ms, TR = 5520 ms, slice thickness = 1.25 mm, flip angle = 78 degrees, FOV = 210*180, voxel size = 1.25mm isotropic). The scans collected at Vanderbilt included healthy controls from several projects (Philips 3T Achieva). A typical acquisition is below, although some variations exist across projects. T1-weighted images acquired using an MPRAGE sequence (TE =2.9 ms, TR = 6.3 ms, slice thickness = 1 mm, flip angle = 8 deg, FOV = 256×240mm, acquisition matrix = 256×240, voxel size = 1×1×1mm). Diffusion images were acquired using a single-shot EPI sequence, and consisted of a single b-value (b = 1000 s/mm^2^), with 65 volumes (1 b0 + 64 DWIs per shell) acquired axially (TE = 101 ms, TR = 5891 ms, slice thickness = 2.2 mm, flip angle = 90 degrees, FOV = 220*220, acquisition matrix = 144*144, voxel size = 2.2mm isotropic).

Data pre-processing included correction for susceptibility distortions, subject motion, eddy current correction[48], and b-table correction[49].

### 2.2 Subject-level processing: tractography

Six methods for tractography and virtual bundle dissection were employed on all diffusion datasets in native space (**Figure 2, Subject-level processing**). These included (1) TractSeg[50], (2) Recobundles[45], (3) Tracula[33], (4) XTract[51], (5) Automatic Fiber-tract Quantification (AFQ)[52], and (6) post-processing of AFQ where only the stem of the bundle was retained, which we call AFQ-clipped.

Algorithms were chosen because they are fully automated, validated, and represent a selection of the state-of-the art methods in the field. In all cases, algorithms were run using default parameters or parameters recommended by original authors.

Briefly, TractSeg is based on convolutional neural networks and performs bundle-specific tractography based on a field of estimated fiber orientations[50, 53], and delineates 72 bundles. We implemented the dockerized version at (https://github.com/MIC-DKFZ/TractSeg). Recobundles segments streamlines based on their shape-similarity to a dictionary of expertly delineated model bundles. Recobundles was run using DIPY[54] software (https://dipy.org) after performing whole-brain tractography. The bundle-dictionary contains 80 bundles, but only 44 were selected to be included in the Pandora atlas after consulting with the algorithm developers based on internal quality assurance (for example removing cranial nerves which are often not used in brain imaging). Of note, Recobundles is a method to automatically extract and recognize bundles of streamlines using prior bundle models, and the implementation we chose uses the DIPY bundle dictionary for extraction, although others can be used. Tracula (https://surfer.nmr.mgh.harvard.edu/fswiki/Tracula) uses probabilistic tractography with anatomical priors based on an atlas and Freesurfer[55–57] (https://surfer.nmr.mgh.harvard.edu) cortical parcellations to constrain the tractography reconstructions. Tracula resulted in 18 bundles segmented per subject. Xtract (https://fsl.fmrib.ox.ac.uk/fsl/fslwiki/XTRACT) is a recent automated method for probabilistic tractography based on carefully selected inclusion, exclusion, and seed regions, selected for 42 tracts in the human brain. AFQ (https://github.com/yeatmanlab/AFQ) is a technique that identifies the core of the major fiber tracts with the aim of quantifying tissue properties within and along the tract, although we only extracted the bundle profile itself. In our case, we extracted the full profile of the bundle, as well as the core of the bundle which was performed in the AFQ software by a clipping operation. For this reason, we called these AFQ and AFQ-clipped, respectively. Both of these methods resulted in 20 bundles. In total, we present 216 bundles in the atlas. A list of the bundles from each pipeline is given in **Appendix A**.

Output from all algorithms were in the form of streamlines, tract-density maps, or probability maps. In all cases, pathways were binarized at the subject level, indicating the voxel-wise existence or non-existence of the bundle in that subject, for that pathway. These binary maps were used to create the population atlases after deformation to standard space.

Exhaustive manual quality assurance (QA) was performed on tractography results. QA included displaying overlays of binarized pathways over select slices for all subjects, inspecting and verifying appropriate shape and location of all bundles on all subjects. We note that not all methods were able to successfully reconstruct all pathways on all subjects, for this reason, some atlases contain information from slightly fewer than all 2443 subjects. Tractography scripts and singularity/dockerized containers as well as QA scripts are provided at (https://github.com/MASILab/Pandora-Methods).

### 2.3 Subject-level processing: registration

In order to create the atlases, all images were registered and transformed to a standard space (**Figure 2, Subject-level processing**). For this work, we chose the MNI standard space, a commonly used space in neuroimaging literature. To do this, the T1 image was intensity normalized using FreeSurfer’s *mri_nu_correct*, *mni*, and *mri_normalize* which perform N3 bias field correction and intensity normalization, respectively on the input T1 image[58]. Next, the diffusion b0 image was coregistered to the T1 using FSL’s *epi_reg[59]* (a rigid-body 6 degrees of freedom transformation). The T1 was then nonlinearly registered using ANTS *antsRegistrationSyn* to a 1.0 mm isotropic MNI ICBM 152 asymmetric template[60]. The FSL transform from *epi_reg* was converted to ANTS format using the *c3d_affine_tool*. Afterwards, all data could be transferred from subject native diffusion space to MNI space (and vice-versa) through *antsApplyTransforms* tools. Thus, all binarized pathways for all subjects were transformed to MNI space using both linear and nonlinear transforms. Transforms were also applied to the normalized T1 images to transform these structural images to standard space.

QA was performed to verify acceptable image registration. This again included generating and visualizing overlays of the b0 images, pathways, and T1 images in MNI spaces overlaid and/or adjacent to the MNI ICBM template image. Both normalization and registration scripts, as well as QA scripts, are provided at (https://github.com/MASILab/Pandora-Methods).

### 2.4 Volumetric atlas creation

Once all data were in MNI space, population-based atlases were created by following methods previously used to create tractography atlases[41, 61, 62]. For each pathway, the binarized maps were summed and set to a probabilistic map between 0 and 100% population overlap (**Figure 2, Volumetric Atlas**). Thus, each pathway was represented as a 3D volume, and concatenation of all volumes results in the 4D volumetric atlas. Atlases were additionally separated based on the method used to create the atlas, as well as separated by dataset (BLSA, HCP, VU) if population-specific or method-specific analysis is required (see *Technical Validation*, below). Scripts for volumetric atlas generation are provided at (https://github.com/MASILab/Pandora-Methods).

### 2.5 Surface-intersection atlas creation

To overlay each pathway onto the MNI template surfaces, a standard FreeSurfer pipeline[58] was used to reconstruct the white/gray matter cortical surfaces directly from the MNI ICBM template image. Each of the probability maps overlaid over the volumetric atlas was then transferred to the reconstructed surfaces to create the surface atlas. However, the reconstructed cortical surfaces do not necessarily guarantee unique voxel-to-vertex matching (normally, more than one vertex belongs to a single voxel) even if they perfectly trace the white- and gray-matter boundary. This potentially degenerates vertex-to-voxel mapping without a voxel-wise resampling scheme. Therefore, the probability to a given vertex was obtained by tri-linear resampling of the associated voxel for sub-voxel accuracy. Scripts for surface atlas generation are provided at (https://github.com/MASILab/Pandora-Methods).

### 2.6 Data visualization and validation

Qualitative validation of the atlases included pathway visualization as an overlay of the population probability on the MNI ICBM template image, or visualization of population-probability on the white matter/gray matter surface. These displays were used in QA during atlas creation, ensuring acceptable probability values, as well as agreement with expected anatomy, shape, and location.

To quantify similarities and differences across pathways and methods, a pathway-correlation measure was used. The pathway-correlation was calculated between two pathways by taking the correlation coefficient of all voxels where either pathway has a probability > 0. This correlation coefficient ranged from −1 to 1, where a value of 1 indicates a perfect correlation of population densities. Thus, this metric measures the coherence between population maps obtained from the bundles and was used to assess if the distribution of population probabilities in space is similar. We used this measure to test similarities/differences between the pathways from different bundle dissection methods (to justify the use of different tractography methods) as well as between pathways generated from the different datasets (to justify making available atlases separated by dataset, as well as understand differences in results based on populations).

Finally, a uniform manifold approximation and projection (UMAP)[63] was used for dimensionality reduction in order to further assess similarities and differences in pathways across methodologies. The UMAP is a general non-linear dimension reduction that is particularly well suited for visualizing high-dimensional datasets.

## 3. Data Records

All data records described in this manuscript are available through both NITRC and Github repositories (https://www.nitrc.org/projects/pandora_atlas; https://github.com/MASILab/Pandora-WhiteMatterAtlas). The data is composed of several file types, including GNU-zipped NifTi files, VTK files, and atlas meta-data CSV files. For each of the six methods used for WM parcellation (where the nomenclature <METHODS> represents “AFQ”, “AFQ-clipped”, “Recobundles”, “TractSeg”, “Tracula”, and “Xtract”) there are three primary file sub-divisions: (1) volumetric atlases, (2) surface atlas, and (3) meta-data information files.

### 3.1 Volumetric Atlas

The WM volumetric atlases created using the nonlinear registration to standard space are NifTi file formats. For each bundle segmentation method, there is one file which corresponds to the atlas composed of all data with the naming convention <METHOD>.nii.gz, and there are three supplementary files which designate the subset of data the atlas is composed of: “BLSA”, “HCP”, and “VU” with the naming convention <METHOD>_<DATASET>.nii.gz. Here <METHOD> and <DATASET> describe the segmentation method and population dataset.

Each WM pathway corresponds to a single 3D volume of the 4D dataset, stored as double-precision floating-point format with values ranging from 0 to 1. For simplicity, and in line with the template used for data normalization, the image matrix is gridded at 1 mm^3^ isotropic resolution, but other resolutions can be calculated given an appropriate interpolation.

### 3.2 Surface-Intersection Atlas

The surface-intersection atlases created for each method are stored as a VTK file type. For each method and for each hemisphere, there is one file which corresponds to the atlas composed of all data with the name convention <HEMISPHERE>_<METHOD>.vtk.gz, and there are three supplementary files which designate the subset of data the atlas is composed of: “BLSA”, “HCP”, and “VU” with the naming convention <HEMISPHERE>_<METHOD>_<DATASET>.vtk.gz. Here <HEMISPHERE>, <METHOD>, and <DATASET> describe the left or right hemisphere (lh or rh), the segmentation method, and the population dataset.

VTK file contains polygonal data with graphics primitives including vertices, edges, and triangle strips defining the polygonal data. Although the surface mesh itself (i.e., vertices and triangles) is the same for all datasets and methods, each VTK file has a separate dataset attribute that consists of scalar values assigned to each vertex. Thus, within each VTK file there is a set of scalars for each WM pathway, as well as a lookup table name as a character string based on the name of the specific pathway.

### 3.3 Meta-data

Each <METHOD> has an associated meta-data file stored as a CSV file. This file contains a numerical index for every volume within the atlas. With every number there is also an associated anatomical label for the pathway, a “file-system” name using a label – typically an acronym – that is friendly for scripting (no spaces or special characters), as well as the number of subjects from each database that are included in the creation of the population-template for that pathway.

### 3.4 Additional data: T1 template, linear atlases, gray matter atlases, and scripting

In addition to the WM labels, a number of supplementary data are also provided with the atlas. First, we created the T1-averaged template derived from all datasets, as well as each dataset separately, and make these available for use as an alternative to the MNI ICBM template, if desired. Second, we also created WM volumetric atlases created using linear registration to MNI space and provide these as supplementary data within each method. Third, we created the population-average of several commonly used gray matter parcellations, including brainCOLOR labels, and also the Desikan-Killany[26], Destrieux[55], and DKTatlas40[64] parcellation schemes from FreeSurfer. Every subjects T1 image was labeled in native space, and labels were warped to standard space where majority voting was applied to produce labels in atlas space. These gray matter labels may provide a reference for localizing the bundles within the WM atlas. Finally, several MATLAB and python scripts are provided to extract individual labels from a given atlas (because many software packages do not facilitate 4D analysis), as well as example normalization and label-propagation scripts.

## 4. Technical Validation

We begin with a qualitative validation of the data, thoroughly inspecting and visualizing all volumes and surfaces from each atlas. An example visualization for 10 selected pathways from the TractSeg sets of atlases is shown in **Figure 3**. All pathways overlay in the correct location, with the correct shape and trajectory, as expected. Population agreement is generally high in the core of the bundle (values ~1) with larger variability along the periphery of pathways. Through this qualitative validation process, differences in the methodologies were noted including some possessing high sensitivity (larger volumes, greater agreement across subjects) and those with higher specificity (smaller, well-defined pathways with lower population agreement).

**Figure 3.**
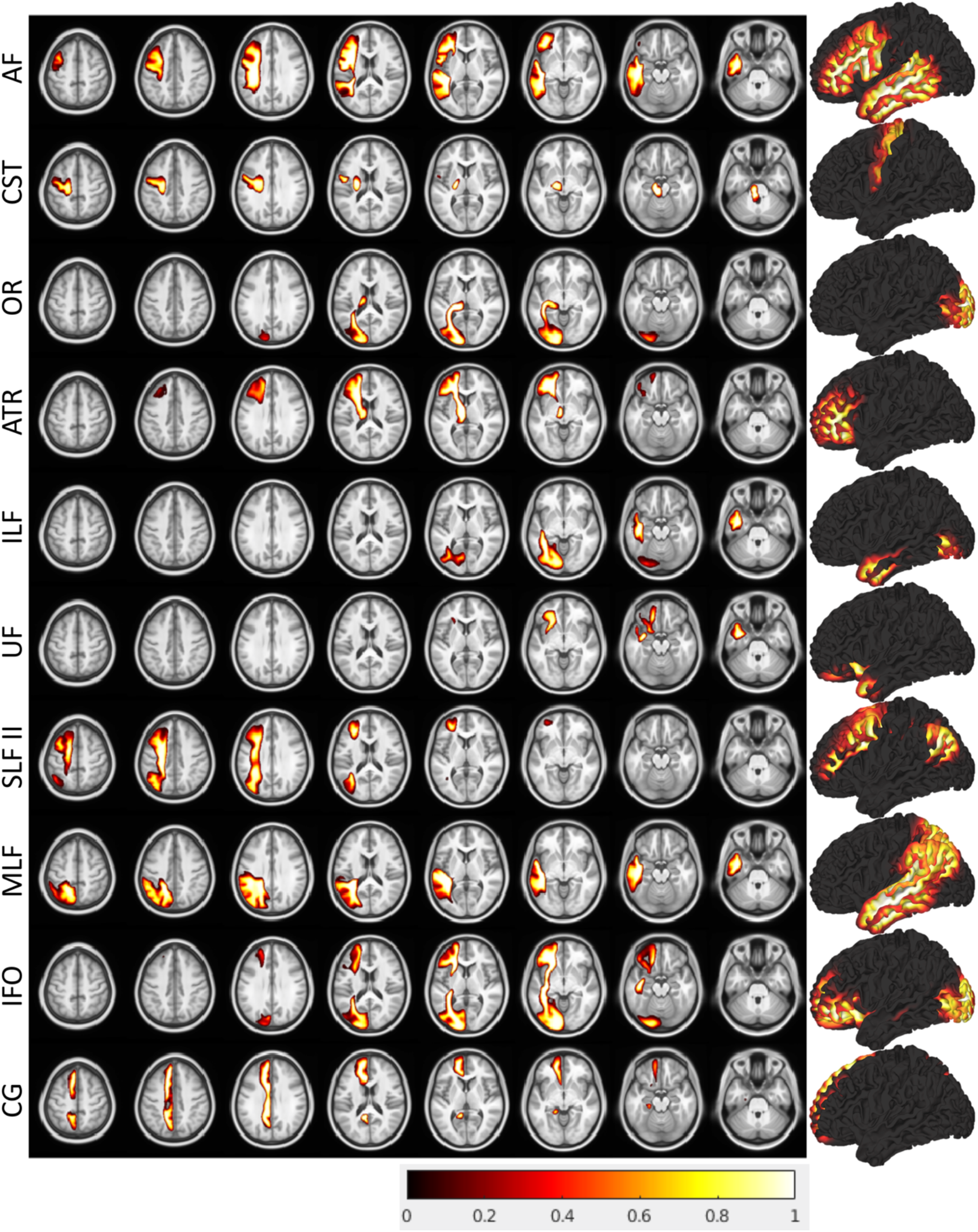
Visualization of data contained in example volumetric and surface atlases. Example visualization for 10 pathways in the TractSeg nonlinear atlas are shown as both overlays and surfaces.

Next, to assess differences within and between tractography techniques, we show pathway-correlations against all other pathways as a large 216×216 matrix of correlations (**Figure 4, a**) and also plotting the UMAP projection of each pathway on a 2D plane (**Figure 4, b**). As expected, most pathways are quite different from others (for example we do not expect the optic radiations to share any overlap whatsoever with the uncinate fasciculus, regardless of methodology), however there are clearly clusters of pathways sharing some similarity, due to both spatial overlap of pathways with comparable anatomies (for example inferior longitudinal fasciculus and inferior frontal occipital fasciculus), as well as methods representing the same pathway. We identified a core group of 20 pathways that are commonly dissected in all methods, and clusters of these pathways are apparent in the UMAP projection (for example, the corticospinal tracts, forceps major and minor, optic radiations, and inferior longitudinal fasciculi are quite similar across algorithms). Thus, *certain pathways are similar, but not exactly the same, across methodologies*, justifying the use of all six state-of-the art methods for bundle dissection.

**Figure 4.**
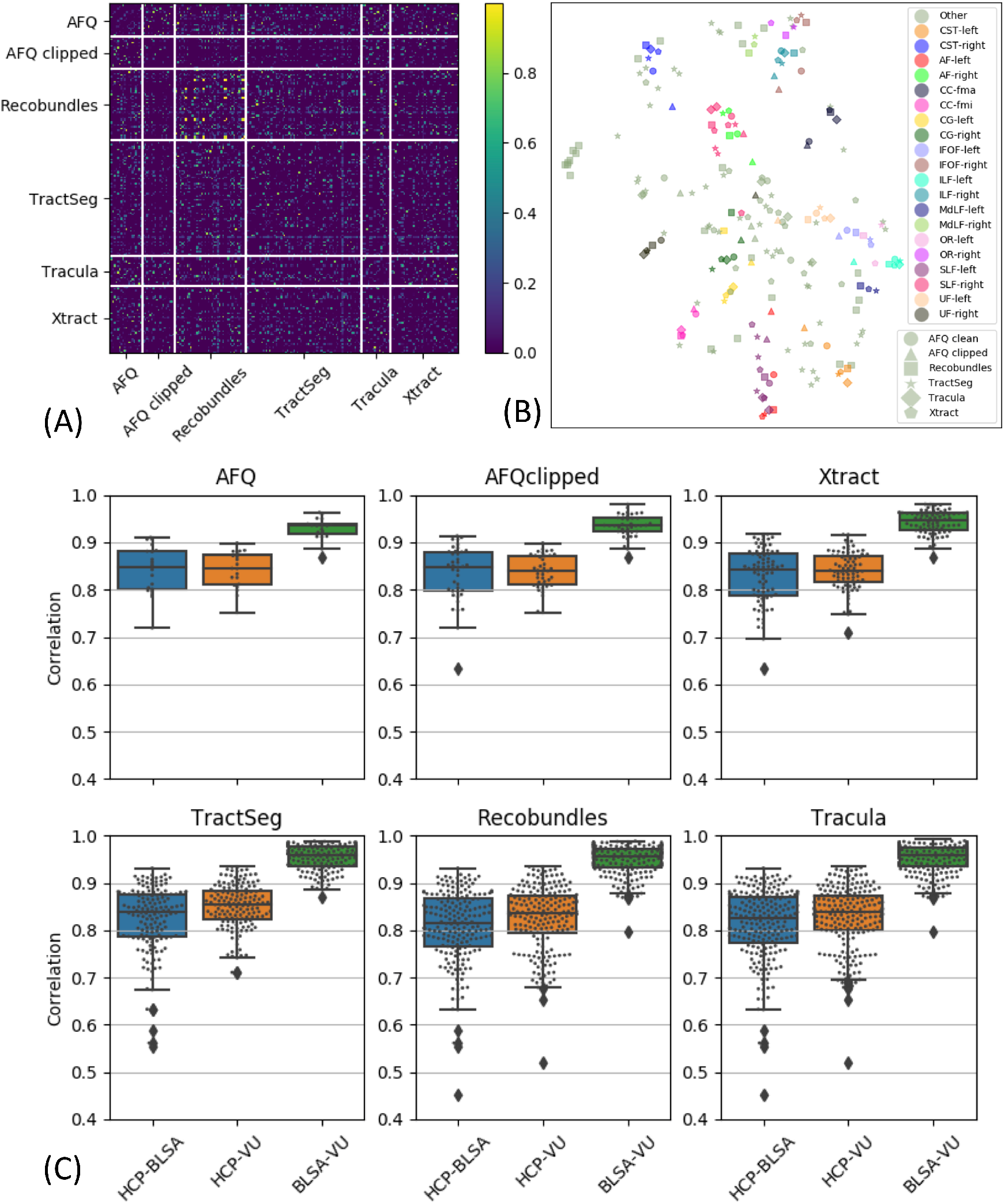
Data validation. (A) Matrix of correlation coefficient of pathways plotted against all others indicates similarities within and across methodologies for bundle dissection. Solid white lines are used to visually separate bundle segmentation methods. (B) UMAP dimensionality reduction projected onto un-scaled 2D plane shows that many WM pathways are similar, but not the same, across methods. Object colors represent specific atlas bundles, with shape indicating segmentation methods. (C) Correlation coefficient of atlases separated by dataset indicates small, but significant, differences between datasets. Together, these justify the inclusion of all tractography methods, as well as separation of atlases by datasets.

Finally, we quantify differences across datasets by showing boxplots of the pathway-correlations after separating by source of data (**Figure 4, c**). While all methods show quite high correlations, it is clear that BLSA and VU datasets and bundles are more similar to each other than to HCP datasets. This is expected as HCP data quality, SNR, resolution, and acquisitions are quite different from the more clinically feasible BLSA and VU sets. Thus, *bundles are also different based on dataset source*. Because of this, in addition to combining results from all subjects, we also supply atlases separated by dataset.

## 5. Usage Notes

Here, we have created and made available the Pandora white matter bundle atlas, that addresses a number of limitations of current human brain atlases by providing a set of population-based volumetric and surface atlases in standard space, based on a large number of subjects, including many pathways from multiple diffusion MRI tractography bundle segmentation methods. We envision the use of these atlases for spatial normalization and label propagation in ways similar to standard usage of volumetric brain atlases. These labels can be used not only for statistical analysis across population and individuals, but also for priors for tractography, relating neuroimaging findings to structural pathways or to inform future methodologies for parcellating and segmenting white matter based on functional, molecular, or alternative contrasts. Similarly, although much less frequently used in the field, the surface-based atlas can also be used to relate functional MRI findings (which are largely applied to cortex, with some evidence for signal contrast in white matter), as priors for cortico-cortical tractography and future bundle segmentations, as a tool for gray matter based spatial statistics, and again for relating alternative neuroimaging findings to structure.

As a simple example workflow. An investigator may be interested in relating tumour localization on a structural image to specific white matter pathways hypothesized to be involved in some functional network. The investigator may choose to register their image to the MNI template, and can either warp their data to template space or apply the inverse transform to get white matter labels into the subject native space. The investigator could then relate tumour location to the probability of given pathways, or could simply threshold the probabilistic maps at a given threshold (for example 0.5) and relate these to the existence/non-existence of the bundle being displaced by the tumour.

We currently recommend the use of the concatenation of all datasets for standard investigative studies unless a population-specific template is required. While differences between datasets are clear and expected, the increased population variability that results from including data from all sources is likely an advantage when investigators are using their own data with possible differences in acquisition, resolution, and subjects. However, future work will investigate creation and dissemination of age-specific white matter analysis, as well as including an age-adjusted surface mesh instead of using the MNI template to generate the surface. We have chosen to include a large number of algorithms for streamline generation and bundle dissection. Our results (**Figure 4**) show that even if the same white matter structure is segmented using different techniques, the results are not guaranteed to be the same. Thus, an investigator could use our atlas with the set of protocols that they agree with most, or alternatively, could relate findings to all white matter pathways across all methodologies in our atlas. We note that we have chosen six standard algorithms to create this atlas, although others exist and new ones are continually developed based on improvements in both our understanding of anatomical connections and our ability to reconstruct these connections with tractography. Inclusion of other tractography and/or segmentation methods are likely additions in future iterations of the atlas, and are easily integrated with existing deformation fields and data organization. Finally, future iterations can include variations and concatenations of gray matter and/or regional atlases in the same space, continually adding to the number of features to be investigated with a single dataset in standard space.

## Code Availability

This atlas is freely available at https://www.nitrc.org/projects/pandora_atlas and https://github.com/MASILab/Pandora-WhiteMatterAtlas. All methods in atlas creation are available at https://github.com/MASILab/Pandora-Methods, including subject-level processing of tractography and registration, volumetric and surface atlas creation, and all QA generation and visualizations.

## Acknowledgments

Research and Education at Vanderbilt University, Nashville, TN. This work was supported by the National Institutes of Health under award numbers R01EB017230 (B.A.L), and T32EB001628 (K.G.S.), R21 MH099218 (W.D.T), R01NS075270 (Victoria L. Morgan) and in part by VICTR VR3029 and the National Center for Research Resources, Grant UL1 RR024975-01, and Department of Defense award number W81XWH-17-2-055. This research was conducted with support from the Intramural Research Program, National Institute on Aging, NIH. The content is solely the responsibility of the authors and does not necessarily represent the official views of the NIH. This work was also supported by the Neuroinformatics Research Chair, Compute Canada, and NSERC Discovery grants of Prof Maxime Descoteaux.

## Author Contributions

CH: conception, investigation, design, analysis, interpretation, writing, revisions; QY: conception, investigation, design, analysis, interpretation, writing, revisions; IL: conception, analysis; FR: conception, design, revisions; CK: investigation, design; BQ: investigation, design; SF: investigation, design; OW: revisions, data; AS: revisions, data; SR: revisions, data; DZ: revisions, data; LC: revisions, data; WT: revisions, data; BB: revisions, data; EG: investigation, design; AA: conception, design; BL: conception, design, analysis, acquisition, interpretation; KS: conception, investigation, design, analysis, interpretation, writing, revisions; MD: conception, design, revisions; KS: conception, investigation, design, analysis, interpretation, writing, revisions

## Appendix A

The bundles resulting from each bundle-segmentation pipeline are given as a list below.

AFQ: Corpus Callosum Forceps Major; Corpus Callosum Forceps Minor; Arcuate Fasciculus left; Cingulum-Cingulate Gyrus left; Cingulum-Hippocampal Gyrus left; Corticospinal Tract left; Inferior Occipito-frontal Fasciculus left; Inferior Longitudinal Fasciculus left; Superior Longitudinal Fasciculus left; Thalamic Radiation left; Uncinate Fasciculus left; Arcuate Fasciculus right; Cingulum-Cingulate Gyrus right; Cingulum-Hippocampal Gyrus right; Corticospinal Tract right; Inferior Occipito-frontal Fasciculus right; Inferior Longitudinal Fasciculus right; Superior Longitudinal Fasciculus right; Thalamic Radiation right; Uncinate Fasciculus right

AFQ-clipped: Corpus Callosum Forceps Major; Corpus Callosum Forceps Minor; Arcuate Fasciculus left; Cingulum-Cingulate Gyrus left; Cingulum-Hippocampal Gyrus left; Corticospinal Tract left; Inferior Occipito-frontal Fasciculus left; Inferior Longitudinal Fasciculus left; Superior Longitudinal Fasciculus left; Thalamic Radiation left; Uncinate Fasciculus left; Arcuate Fasciculus right; Cingulum-Cingulate Gyrus right; Cingulum-Hippocampal Gyrus right; Corticospinal Tract right; Inferior Occipito-frontal Fasciculus right; Inferior Longitudinal Fasciculus right; Superior Longitudinal Fasciculus right; Thalamic Radiation right; Uncinate Fasciculus right

Recobundles: Arcuate Fasciculus left; Arcuate Fasciculus left; Frontal Aslant Tract left; Frontal Aslant Tract right; Cerebellum left; Cerebellum right; Corpus Callosum Major; Corpus Callosum Minor; Central Tegmental Tract left; Central Tegmental Tract right; Extreme Capsule left; Extreme Capsule right; Fronto-pontine tract left; Fronto-pontine tract right; Inferior Fronto-occipital Fasciculus left; Inferior Fronto-occipital Fasciculus right; Inferior Longitudinal Fasciculus left; Inferior Longitudinal Fasciculus right; Middle Cerebellar Peduncle; Middle Longitudinal Fasciculus left; Middle Longitudinal Fasciculus right; Medial Longitudinal fasciculus left; Medial Longitudinal fasciculus right; Medial Lemniscus left; Medial Lemniscus right; Occipito Pontine Tract left; Occipito Pontine Tract right; Optic Radiation left; Optic Radiation right; Parieto Pontine Tract left; Parieto Pontine Tract right; Superior longitudinal fasciculus left; Superior longitudinal fasciculus right; Spinothalamic Tract left; Spinothalamic Tract right; Temporopontine Tract left; Temporopontine Tract right; Uncinate Fasciculus left; Uncinate Fasciculus right; Vermis

TractSeg: Arcuate fascicle left; Arcuate fascicle right; Anterior Thalamic Radiation left; Thalamic Radiation right; Commissure Anterior; Rostrum; Genu; Rostral body (Premotor); Anterior midbody (Primary Motor); Posterior midbody (Primary Somatosensory); Isthmus; Splenium; Corpus Callosum – all; Cingulum left; Cingulum right; Corticospinal tract left; Corticospinal tract right; Fronto-pontine tract left; Fronto-pontine tract right; Fornix left; Fornix right; Inferior cerebellar peduncle left; Inferior cerebellar peduncle right; Inferior occipito-frontal fascicle left; Inferior occipito-frontal fascicle right; Inferior longitudinal fascicle left; Inferior longitudinal fascicle right; Middle cerebellar peduncle; Middle longitudinal fascicle left; Middle longitudinal fascicle right; Optic radiation left; Optic radiation right; Parieto-occipital pontine left; Parieto-occipital pontine right; Superior cerebellar peduncle left; Superior cerebellar peduncle right; Superior longitudinal fascicle III left; Superior longitudinal fascicle III right; Superior longitudinal fascicle II left; Superior longitudinal fascicle II right; Superior longitudinal fascicle I left; Superior longitudinal fascicle I right;Striato-fronto-orbital left; Striato-fronto-orbital right; Striato-occipital left; Striato-occipital right; Striato-parietal left; Striato-parietal right; Striato-postcentral left; Striato-postcentral right; Striato-precentral left; Striato-precentral right; Striato-prefrontal left; Striato-prefrontal right; Striato-premotor left; Striato-premotor right; Superior Thalamic Radiation left; Superior Thalamic Radiation right; Thalamo-occipital left; Thalamo-occipital right; Thalamo-parietal left; Thalamo-parietal right; Thalamo-postcentral left; Thalamo-postcentral right; Thalamo-precentral left; Thalamo-precentral right; Thalamo-prefrontal left; Thalamo-prefrontal right; Thalamo-premotor left; Thalamo-premotor right; Uncinate fascicle left; Uncinate fascicle right

Tracula: Corpus Callosum Forceps Major; Corpus Callosum Forceps Minor; Anterior Thalamic Radiation left; Cingulum - Angular Bundle left; Cingulum - Cingulate Gyrus left; Corticospinal Tract left; Inferior Longitudinal Fasciculus left; Superior Longitudinal Fasciculus - Parietal left; Superior Longitudinal Fasciculus - Temporal left; Uncinate Fasciculus left; Anterior Thalamic Radiation right; Cingulum - Angular Bundle right; Cingulum - Cingulate Gyrus right; Corticospinal Tract right; Inferior Longitudinal Fasciculus right; Superior Longitudinal Fasciculus - Parietal right; Superior Longitudinal Fasciculus - Temporal right; Uncinate Fasciculus right

Xtract: Anterior Commissure; Arcuate Fascile left; Arcuate Fascile right; Acoustic Radiation left; Acoustic Radiation right; Anterior Thalamic Radiation left; Anterior Thalamic Radiation right; Cingulum Bundle Dorsal left; Cingulum Bundle Dorsal right; Cingulum Bundle Parahippocampal left; Cingulum Bundle Parahippocampal right; Cingulum Bundle Temporal left; Cingulum Bundle Temporal right; Corticospinal Tract left; Corticospinal Tract right; Frontal Aslant left; Frontal Aslant right; Forceps Major; Forceps Minor; Fornix left; Fornix right; Inferior Fronto-occipital Fasciculus left; Inferior Fronto-occipital Fasciculus right; Inferior Longitudinal Fasciculus left; Inferior Longitudinal Fasciculus right; Middle Cerebellar Peduncle; Medio-Dorsal Longitudinal Fasciculus left; Medio-Dorsal Longitudinal Fasciculus right; Optic Radiation left; Optic Radiation right; Superior Longitudinal Fasciculus 1 left; Superior Longitudinal Fasciculus 1 right; Superior Longitudinal Fasciculus 2 left; Superior Longitudinal Fasciculus 2 right; Superior Longitudinal Fasciculus 3 left; Superior Longitudinal Fasciculus 3 right; Superior Thalamic Radiation left; Superior Thalamic Radiation right; Uncinate Fasciculus left; Uncinate Fasciculus right; Vertical Occipital Fasciculus left; Vertical Occipital Fasciculus right;

## Footnotes

* This name was chosen as a parallel to the JHU “Eve” atlas, where Pandora was the first woman in Greek mythology. Also, from Pandora’s box was released “evil” and only “hope” remained. Our Pandora’s “box” just happens to contain white matter labels.

## References

1. Cabezas, M., et al., A review of atlas-based segmentation for magnetic resonance brain images. Comput Methods Programs Biomed, 2011. 104(3): p. e158–77.

2. Toga, A.W., Brain warping. 1999, San Diego: Academic Press. xiii, 385 p.

3. Gee, J.C., M. Reivich, and R. Bajcsy, Elastically deforming 3D atlas to match anatomical brain images. J Comput Assist Tomogr, 1993. 17(2): p. 225–36.

4. Myers, P.E., et al., Standardizing Human Brain Parcellations. bioRxiv, 2019: p. 845065.

5. Yeo, B.T., et al., The organization of the human cerebral cortex estimated by intrinsic functional connectivity. J Neurophysiol, 2011. 106(3): p. 1125–65.

6. Schaefer, A., et al., Local-Global Parcellation of the Human Cerebral Cortex from Intrinsic Functional Connectivity MRI. Cereb Cortex, 2018. 28(9): p. 3095–3114.

7. Glasser, M.F., et al., A multi-modal parcellation of human cerebral cortex. Nature, 2016. 536(7615): p. 171–178.

8. Rolls, E.T., et al., Automated anatomical labelling atlas 3. Neuroimage, 2020. 206: p. 116189.

9. Tzourio-Mazoyer, N., et al., Automated anatomical labeling of activations in SPM using a macroscopic anatomical parcellation of the MNI MRI single-subject brain. Neuroimage, 2002. 15(1): p. 273–89.

10. Mars, R.B., et al., Diffusion-weighted imaging tractography-based parcellation of the human parietal cortex and comparison with human and macaque resting-state functional connectivity. J Neurosci, 2011. 31(11): p. 4087–100.

11. Mars, R.B., et al., Connectivity-based subdivisions of the human right “temporoparietal junction area”: evidence for different areas participating in different cortical networks. Cereb Cortex, 2012. 22(8): p. 1894–903.

12. Sallet, J., et al., The organization of dorsal frontal cortex in humans and macaques. J Neurosci, 2013. 33(30): p. 12255–74.

13. Neubert, F.X., et al., Comparison of human ventral frontal cortex areas for cognitive control and language with areas in monkey frontal cortex. Neuron, 2014. 81(3): p. 700–13.

14. Neubert, F.X., et al., Connectivity reveals relationship of brain areas for reward-guided learning and decision making in human and monkey frontal cortex. Proc Natl Acad Sci U S A, 2015. 112(20): p. E2695–704.

15. Labache, L., et al., A SENtence Supramodal Areas AtlaS (SENSAAS) based on multiple task-induced activation mapping and graph analysis of intrinsic connectivity in 144 healthy right-handers. Brain Struct Funct, 2019. 224(2): p. 859–882.

16. Kikinis, R., et al., A digital brain atlas for surgical planning, model-driven segmentation, and teaching. IEEE Transactions on Visualization and Computer Graphics, 1996. 2(3): p. 232–241.

17. van Baarsen, K.M., et al., A probabilistic atlas of the cerebellar white matter. Neuroimage, 2015. 124(Pt A): p. 724–732.

18. Mazziotta, J.C., et al., A probabilistic atlas of the human brain: theory and rationale for its development. The International Consortium for Brain Mapping (ICBM). Neuroimage, 1995. 2(2): p. 89–101.

19. Behrens, T.E., et al., Non-invasive mapping of connections between human thalamus and cortex using diffusion imaging. Nat Neurosci, 2003. 6(7): p. 750–7.

20. Ewert, S., et al., Toward defining deep brain stimulation targets in MNI space: A subcortical atlas based on multimodal MRI, histology and structural connectivity. Neuroimage, 2018. 170: p. 271–282.

21. Ilinsky, I., et al., Human Motor Thalamus Reconstructed in 3D from Continuous Sagittal Sections with Identified Subcortical Afferent Territories. eNeuro, 2018. 5(3).

22. Keuken, M.C., et al., Ultra-high 7T MRI of structural age-related changes of the subthalamic nucleus. J Neurosci, 2013. 33(11): p. 4896–900.

23. Tziortzi, A.C., et al., Imaging dopamine receptors in humans with [11C]-(+)-PHNO: dissection of D3 signal and anatomy. Neuroimage, 2011. 54(1): p. 264–77.

24. Mazziotta, J., et al., A probabilistic atlas and reference system for the human brain: International Consortium for Brain Mapping (ICBM). Philos Trans R Soc Lond B Biol Sci, 2001. 356(1412): p. 1293–322.

25. Makris, N., et al., Decreased volume of left and total anterior insular lobule in schizophrenia. Schizophr Res, 2006. 83(2-3): p. 155–71.

26. Desikan, R.S., et al., An automated labeling system for subdividing the human cerebral cortex on MRI scans into gyral based regions of interest. Neuroimage, 2006. 31(3): p. 968–80.

27. Eickhoff, S.B., et al., Assignment of functional activations to probabilistic cytoarchitectonic areas revisited. Neuroimage, 2007. 36(3): p. 511–21.

28. Eickhoff, S.B., et al., Testing anatomically specified hypotheses in functional imaging using cytoarchitectonic maps. Neuroimage, 2006. 32(2): p. 570–82.

29. Eickhoff, S.B., et al., A new SPM toolbox for combining probabilistic cytoarchitectonic maps and functional imaging data. Neuroimage, 2005. 25(4): p. 1325–35.

30. Talairach, J. and P. Tournoux, Co-planar stereotaxic atlas of the human brain : 3-dimensional proportional system : an approach to cerebral imaging. 1988, Stuttgart; New York: Georg Thieme. 122 p.

31. Lancaster, J.L., et al., Automated Talairach atlas labels for functional brain mapping. Hum Brain Mapp, 2000. 10(3): p. 120–31.

32. Oishi, K., et al., Atlas-based whole brain white matter analysis using large deformation diffeomorphic metric mapping: application to normal elderly and Alzheimer’s disease participants. Neuroimage, 2009. 46(2): p. 486–99.

33. Yendiki, A., et al., Automated probabilistic reconstruction of white-matter pathways in health and disease using an atlas of the underlying anatomy. Front Neuroinform, 2011. 5: p. 23.

34. Mori, S., et al., Stereotaxic white matter atlas based on diffusion tensor imaging in an ICBM template. Neuroimage, 2008. 40(2): p. 570–82.

35. Mori, S., et al., MRI Atlas of Human White Matter. 2005: Academic Press, 2010. 276.

36. Catani, M. and M. Thiebaut de Schotten, A diffusion tensor imaging tractography atlas for virtual in vivo dissections. Cortex, 2008. 44(8): p. 1105–32.

37. Oishi, K., et al., Human brain white matter atlas: identification and assignment of common anatomical structures in superficial white matter. Neuroimage, 2008. 43(3): p. 447–57.

38. Roman, C., et al., Clustering of Whole-Brain White Matter Short Association Bundles Using HARDI Data. Front Neuroinform, 2017. 11: p. 73.

39. Yeh, F.C., et al., Population-averaged atlas of the macroscale human structural connectome and its network topology. Neuroimage, 2018. 178: p. 57–68.

40. Yeh, F.C. and W.Y. Tseng, NTU-90: a high angular resolution brain atlas constructed by q-space diffeomorphic reconstruction. Neuroimage, 2011. 58(1): p. 91–9.

41. Chenot, Q., et al., A population-based atlas of the human pyramidal tract in 410 healthy participants. Brain Struct Funct, 2019. 224(2): p. 599–612.

42. Archer, D.B., D.E. Vaillancourt, and S.A. Coombes, A Template and Probabilistic Atlas of the Human Sensorimotor Tracts using Diffusion MRI. Cereb Cortex, 2018. 28(5): p. 1685–1699.

43. Rojkova, K., et al., Atlasing the frontal lobe connections and their variability due to age and education: a spherical deconvolution tractography study. Brain Structure and Function, 2016. 221(3): p. 1751–1766.

44. Thiebaut de Schotten, M., et al., Monkey to human comparative anatomy of the frontal lobe association tracts. Cortex, 2012. 48(1): p. 82–96.

45. Garyfallidis, E., et al., Recognition of white matter bundles using local and global streamline-based registration and clustering. Neuroimage, 2018. 170: p. 283–295.

46. Zhang, F., et al., An anatomically curated fiber clustering white matter atlas for consistent white matter tract parcellation across the lifespan. Neuroimage, 2018. 179: p. 429–447.

47. Guevara, P., et al., Automatic fiber bundle segmentation in massive tractography datasets using a multi-subject bundle atlas. Neuroimage, 2012. 61(4): p. 1083–99.

48. Andersson, J.L., S. Skare, and J. Ashburner, How to correct susceptibility distortions in spin-echo echo-planar images: application to diffusion tensor imaging. Neuroimage, 2003. 20(2): p. 870–88.

49. Schilling, K.G., et al., A fiber coherence index for quality control of B-table orientation in diffusion MRI scans. Magn Reson Imaging, 2019. 58: p. 82–89.

50. Wasserthal, J., P. Neher, and K.H. Maier-Hein, TractSeg - Fast and accurate white matter tract segmentation. Neuroimage, 2018. 183: p. 239–253.

51. Warrington, S., et al., XTRACT - Standardised protocols for automated tractography and connectivity blueprints in the human and macaque brain. bioRxiv, 2019: p. 804641.

52. Yeatman, J.D., et al., Tract profiles of white matter properties: automating fiber-tract quantification. PLoS One, 2012. 7(11): p. e49790.

53. Wasserthal, J., et al., Combined tract segmentation and orientation mapping for bundle-specific tractography. Med Image Anal, 2019. 58: p. 101559.

54. Garyfallidis, E., et al., Dipy, a library for the analysis of diffusion MRI data. Front Neuroinform, 2014. 8: p. 8.

55. Destrieux, C., et al., Automatic parcellation of human cortical gyri and sulci using standard anatomical nomenclature. Neuroimage, 2010. 53(1): p. 1–15.

56. Fischl, B., M.I. Sereno, and A.M. Dale, Cortical surface-based analysis. II: Inflation, flattening, and a surface-based coordinate system. Neuroimage, 1999. 9(2): p. 195–207.

57. Dale, A.M., B. Fischl, and M.I. Sereno, Cortical surface-based analysis. I. Segmentation and surface reconstruction. Neuroimage, 1999. 9(2): p. 179–94.

58. Fischl, B., FreeSurfer. Neuroimage, 2012. 62(2): p. 774–781.

59. Smith, S.M., et al., Advances in functional and structural MR image analysis and implementation as FSL. Neuroimage, 2004. 23: p. S208–S219.

60. Avants, B.B., N. Tustison, and G. Song, Advanced normalization tools (ANTS). Insight j, 2009. 2: p. 1–35.

61. Bürgel, U., et al., White matter fiber tracts of the human brain: Three-dimensional mapping at microscopic resolution, topography and intersubject variability. NeuroImage, 2006. 29(4): p. 1092–1105.

62. Thiebaut de Schotten, M., et al., Atlasing location, asymmetry and inter-subject variability of white matter tracts in the human brain with MR diffusion tractography. Neuroimage, 2011. 54(1): p. 49–59.

63. Melville, L.M.a.J.H.a.J., UMAP: Uniform Manifold Approximation and Projection for Dimension Reduction. 2018, arXiv: 1802.03426.

64. Klein, A. and J. Tourville, 101 labeled brain images and a consistent human cortical labeling protocol. Front Neurosci, 2012. 6: p. 171.

